# Temperature but not ocean acidification affects energy metabolism and enzyme activities in the blue mussel, *Mytilus edulis*

**DOI:** 10.1101/2020.11.29.402933

**Authors:** Omera B. Matoo, Gisela Lannig, Christian Bock, Inna M. Sokolova

## Abstract

1. In mosaic marine habitats such as intertidal zones ocean acidification (OA) is exacerbated by high variability of pH, temperature, and biological CO_2_ production. The non-linear interactions among these drivers can be context-specific and their effect on organisms in these habitats remains largely unknown, warranting further investigation.
2. We were particularly interested in *Mytilus edulis* (the blue mussel) from intertidal zones of Gulf of Maine (GOM), USA for this study. GOM is a hot spot of global climate change (average SST increasing by > 0.2 °C y^-1^) with > 60% decline in mussel population over the past 40 years.
3. Here, we utilize bioenergetic underpinnings to identify limits of stress tolerance in *M. edulis* from GOM exposed to warming and OA. We have measured whole-organism oxygen consumption rates and metabolic biomarkers in mussels exposed to control and elevated temperatures (10 vs. 15 °C) and moderate P_CO2_ levels (~ 400 vs. 800 μatm).
4. Our study demonstrates that adult *M. edulis* from GOM are metabolically resilient to the moderate OA scenario but responsive to warming as seen in changes in metabolic rate, energy reserves, metabolite profiles and enzyme activities.
5. Our results are in agreement with recent literature that OA scenarios for the next 100-300 years do not affect this species, possibly as a consequence of maintaining its *in vivo* acid-base balance.

## INTRODUCTION

Continued increase in atmospheric CO_2_ and its subsequent uptake by oceans profoundly affects marine ecosystems (IPCC, 2014). Changes experienced by organisms include increase in global sea surface temperature (SST) and oceans’ partial pressure of CO_2_ (P_CO2_), which leads to ocean acidification (OA) (Doney *et al*., 2012). Climate change models predict an average increase of 1.8–4.0 °C (with some estimates as high as 6.4 °C) and a decline by 0.3-0.4 pH units by the year 2100, depending on the CO_2_ emission scenario (IPCC, 2014). Warming and OA can negatively impact marine organisms (Kroeker *et al*., 2013, 2014). However, in mosaic habitats such as intertidal and coastal zones, the outcome of multiple drivers, including warming and OA, is complicated by context-specific and non-linear interactions among the drivers (Gunderson *et al*., 2016 and references therein) so that the net effect could be additive, antagonistic or synergistic (Todgham & Stillman, 2013). The species’ response to interactive effects of warming and OA in such environments remains largely unknown and warrants further investigation (Gunderson *et al*., 2016).

Temperature is a key variable that affects physiology, survival, and distribution of ectotherms (Kroeker *et al*., 2013, 2014). Deviation of temperature from the optimum results in disturbance of energy balance and decrease in aerobic scope of organisms (Pörtner, 2012; Sokolova, 2013; Sokolova *et al*., 2012). OA negatively affects survival, metabolism, calcification, growth, reproduction and immune responses across a range of marine taxa (Kroeker *et al*., 2013). Elevated P_CO2_ shifts the acid-base balance of organisms (Melzner *et al*., 2009) and most calcifiers have limited capacity to counteract OA-induced extracellular acidosis (Pörtner, 2008). This in turn can increase the energy costs to maintain cellular and organismal homeostasis in animals (Ivanina *et al*., 2020; Pörtner, 2008; Sokolova *et al*., 2012; Stapp *et al*., 2018; Stumpp *et al*., 2012).

Responses of marine molluscs to OA are highly variable (Sokolova *et al*., 2015 and references therein). Inter- and intrapopulation variability in OA sensitivity has been shown depending on habitat, scales of environmental variability and other concomitant stressors (Vargas *et al*., 2017; Waldbusser *et al*., 2015; Parker *et al*., 2011; Stapp *et al*., 2017). In mosaic environments local adaptation as well as temporally and spatially varying selection can select for metabolically plastic, stress-tolerant genotypes that can maintain optimal phenotypes (including energetic sustainability) in a broad range of environmental conditions. Importantly, bioenergetic responses can predict tolerance limits under environmentally realistic scenarios of stress exposure (Sokolova *et al*., 2012) providing a common denominator to integrate responses to multiple stressors. Quantifying the independent and interactive effects of multiple stressors to identify metabolic tipping points is essential to determine the impact of global climate change on marine organisms and ecosystems (Boyd & Brown, 2015)

This study aims to determine the interactive effects of elevated temperature and P_CO2_ on energy metabolism and biomineralization-related enzymes in an ecologically and economically important bivalve mollusk, the blue mussel *Mytilus edulis* Linnaeus 1758. It is a critical foundation species in coastal ecosystems (Seed, 1969) that increasingly faces risk of local extinction along the USA east coast (Jones *et al*., 2009; Sorte *et al*., 2017). We were particularly interested in mussel populations from intertidal zones of Gulf of Maine (GOM), USA for this study. GOM is a hot spot of global climate change with > 60% (range 29-100%) decline in mussel population over the past 40 years (Sorte *et al*., 2017). Within GOM, the previous decade has witnessed SST exceeding earlier recorded observations of >150 years and an average warming of > 0.2 °C y^-1^ (Sorte *et al*., 2017). Mussels inhabiting GOM are thus exposed to one of the fastest rates of warming in the world, in addition to ocean acidification.

The metabolic plasticity of *M. edulis* to the combined effects of elevated P_CO2_ and temperature from GOM is not yet fully understood. Here, we utilize the bioenergetic underpinnings of stress physiology to identify limits and mechanisms of stress tolerance in *M.edulis*. We have measured whole-organism oxygen consumption rates and metabolic biomarkers in mussels exposed to control and moderately elevated temperatures (10 vs. 15 °C) and P_CO2_ (~ 400 vs. 800 μatm) to mimic a realistic scenario of ocean warming and acidification. Standard metabolic rate (SMR) represents the basal energy cost for maintenance and is widely used to assess stress response (Pettersen *et al*., 2018). To account for possible tissue-specific variation in responses, we conducted a comprehensive analyses of the bioenergetic health index by measuring energy related biomarkers (cellular and tissue energy reserves), metabolic profiles and specific enzyme activities (acid-base regulating enzyme and energy-demanding ion transport enzymes) in different tissues depending on their physiological role.

## MATERIALS AND METHODS

### Animal maintenance

*M.edulis* were collected from Biddeford Pool, GOM (43°26’50.6N, 70°21’19.0W) in summer 2011 and shipped overnight to the University of North Carolina at Charlotte. Mussels were kept in recirculating artificial seawater (ASW) (Instant Ocean^®^, USA) at 9.6 ± 0.3 °C and 30 ± 1 salinity (practical salinity units), aerated with ambient air for 10 days. Mussels were randomly assigned to four treatment groups and exposed for four weeks to a combination of two P_CO2_ levels representative of present-day conditions (~400 μatm P_CO2_; normocapnia) and year 2100 projections (~ 800 μatm P_CO2_; hypercapnia) (IPCC, 2014), and two temperatures representing the average water temperature at the time of collection (10 °C), and + 5 °C increase predicted for the year 2100 by IPCC (15 °C). Two replicate tanks were set for each treatment. Normocapnic treatments were bubbled with ambient air whereas for hypercapnia ambient air was mixed with 100% CO_2_ (Roberts Oxygen, USA) using precision mass flow controllers (Cole-Parmer, USA) to maintain a steady-state pH. Animals were fed *ad libitum* on alternate days with 2 mL per tank of commercial algal mixture (Shellfish Diet 1800, USA). Mortality was checked daily. ASW were prepared using the same batch of Instant Ocean^®^ salt to avoid potential variations in water chemistry. Carbonate chemistry of seawater (Table 1) was determined as described elsewhere (Beniash *et al*., 2010).

**Table 1:** Summary of water chemistry parameters during experimental exposure. Salinity, temperature, pH_NBS_ and dissolved inorganic carbon (DIC) was measured in water samples collected during the exposure. Average DIC was 2953.75±111.28 μmol kg^-1^ SW. Other parameters are calculated using co2SYS. Data is represented as Means ± SEM. Same batch of sea water was used throughout the course of the experiment with an average total alkalinity (TA) of 3098.40 mmol kg^-1^ SW.

| Parameters | Assay Temperature |  |  |  |
| --- | --- | --- | --- | --- |
|  | 10 °C |  | 15 °C |  |
|  | Normocapnia | Hypercapnia | Normocapnia | Hypercapnia |
| Salinity, PSU | 30.04±0.07 | 30.50±0.03 | 30.15±0.04 | 30.14±0.05 |
| Temperature, °C | 9.51±0.06 | 9.76±0.02 | 14.95±0.01 | 14.99±0.01 |
| $\text{pH}_{\text{NBS}}$ | 8.13±0.00 | 7.91±0.00 | 8.18±0.00 | 7.92±0.00 |
| $\text{P}_{\text{CO}_2}$, $\mu\text{atm}$ | 585.39±39 | 1026.97±15.07 | 536.14±4.35 | 1069.41±13.90 |
| $\text{HCO}_3^-$, ($\mu\text{mol kg}^{-1} \text{ SW}$) | 2735.82±2.91 | 2863.82±3.49 | 2628.49±2.73 | 2823.22±2.89 |
| $\text{CO}_3^{2-}$, ($\mu\text{mol kg}^{-1} \text{ SW}$) | 155.50±1.23 | 100.37±1.49 | 202.67±1.18 | 118.56±1.24 |
| $\text{CO}_2$, ($\mu\text{mol kg}^{-1} \text{ SW}$) | 26.85±0.26 | 46.42±0.67 | 20.66±0.16 | 41.03±0.53 |
| $\Omega_{\text{calcite}}$ | 3.80±0.02 | 2.44±0.03 | 4.97±0.02 | 2.91±0.03 |
| $\Omega_{\text{aragonite}}$ | 2.38±0.01 | 1.54±0.02 | 3.15±0.01 | 1.84±0.01 |

N = 5 for DIC and N = 36-77 for other parameters.

### Standard Metabolic Rate

SMR was measured as resting oxygen consumption (M O2) of mussels at their respective acclimation temperature and P_CO2_ using microfiber optic oxygen probes (Tx-Type, PreSens GmbH, Germany) as described in Matoo *et al*., 2013. After measurements, individuals were dissected and dry tissue mass was calculated from the wet tissue mass assuming an average water content of 80%.

SMR was calculated as follows:

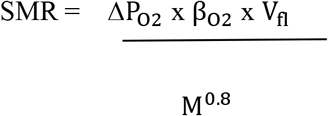

SMR —oxygen consumption (μmol O_2_ g^-1^ dry mass h^-1^) normalized to 1 g dry mass, ΔPO_2_ — difference in partial pressure between in- and out-flowing water (kPa), β_O2_ — oxygen capacity of water (μmol O_2_ L^-1^ kPa^-1^), Vfl — flow rate (L h^-1^), M —dry tissue mass (g) and 0.8 — allometric coefficient (Bougrier et al., 1995).

After acclimation, a subset of mussels was dissected, tissues shock-frozen and stored in liquid nitrogen for analyses of energy reserves and enzyme activities. Due to limited amount of tissues, we divided samples for different assays depending on the physiological function of a given tissue. The energy reserves (lipids, glycogen and/or adenylates) were measured in hepatopancreas (HP) and adductor muscle that serve as reserve storage sites in bivalves (Cappello *et al*., 2018). Metabolite profiles were explored by untargeted metabolomics in two metabolically active aerobic tissues, the gills and the muscle. The effects of warming and OA on biomineralization were assessed by the activities of three key enzymes involved in shell formation (carbonic anhydrase (CA), plasma membrane calcium (Ca^2+^) ATPase and proton (H^+^) ATPase) in the mantle tissue as the main organ involved in shell formation.

### Energy reserves

Lipid content was determined using the chloroform extraction method as described in Ivanina *et al*., 2013. Concentration of lipids were expressed as g g^-1^ wet tissue mass. Concentrations of glycogen and adenylates (μmol g^-1^ wet tissue mass) were measured using standard NADH- or NADPH-linked spectrophotometric tests described in Ivanina *et al*., 2013. Adenylate energy charge (AEC) was calculated as follows:

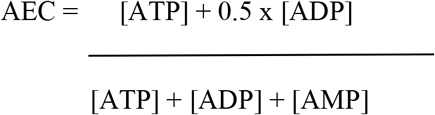

### Metabolic profiling based on ^1^H-NMR spectroscopy

Muscle and gill tissues were extracted as described elsewhere for untargeted metabolic profiling (Dickinson *et al*., 2012; Lannig *et al*., 2010). Extracts were freeze-dried and shipped on dry ice to the Alfred Wegener Institute (Bremerhaven, Germany). Untargeted metabolic profiling using ^1^H-NMR spectroscopy was performed using a method modified from Schmidt *et al*., 2017. 24 metabolites were identified from processed spectra of adductor and gill tissues and quantified using Chenomx NMR suite 8.1 (Chenomx Inc. Canada). Metabolic profiles were statistically analyzed using MetaboAnalyst 4.0 (Chong *et al*. 2019).

### Enzyme activities

Activities of biomineralization-related enzymes were measured in the mantle edge which is involved in shell deposition (Bjärnmark *et al*., 2016; Gazeau *et al*., 2014)] of *M.edulis*. Protein concentrations were determined using Bradford assay (Bradford, 1976) and used to standardize enzyme activities.

#### a) Carbonic anhydrase (CA)

For assessment of carbonic anhydrase (carbonate hydrolyase, EC 4.2.1.1) activity, mantle edge tissue was homogenized as described in Ivanina *et al*., 2013. CA activity was determined as acetazolamide (AZM)-sensitive esterase activity with 1.5 mM of p-nitrophenyl acetate as a substrate (Gambhir *et al*., 2007) using a temperature-controlled spectrophotometer (VARIAN Cary 50 Bio UV–Vis spectrophotometer, USA). In a separate set of experiments, CA activity was measured at different temperatures in an environmentally relevant range (5-35 °C) in the gill, mantle, adductor muscle and HP of the control mussels to characterize the tissue-dependent capacity and temperature sensitivity (determined by apparent activation energy (E_a_) and Arrhenius breakpoint temperature (ABT)) of CA activity. E_a_ was determined from an Arrhenius plot of ln(V_max_) against 1/T (K^-1^), and ABT was determined as a point when the slope of Arrhenius plot significantly changed using an algorithm for multi-segment linear regression (Oosterbaan, 2011).

#### b) Ca^2+^- and H^+^-ATPases

Mantle edge tissue was homogenized and activities of Ca^2+^-ATPase (EC 3.6.3.8) and H^+^-ATPase (EC 3.6.3.6) was assayed as described in Ivanina *et al*., 2020. Inorganic phosphate (Pi) was measured using malachite green assay kit (ab65622, Abcam) and ATPase activities were expressed as μmol of Pi μg protein^-1^ hr^-1^.

### Statistical analyses

Effects of temperature, P_CO2_ and their interaction were assessed for all studied traits using generalized linear model (GLM) ANOVA. All factors were treated as fixed and post-hoc tests (Fisher’s Least Significant Difference) were used to test differences between group means. Sample sizes were 5–10 for all experimental groups. Regression analysis for ABT and Arrhenius plots for CA were done using GraphPad Prism ver. 4.03 (GraphPad Software, Inc.) and SegReg software (Oosterbaan, 2011). For statistical analysis of the metabolic profiles we used MetaboAnalyst 4.0 (Chong *et al*., 2019) as described elsewhere (Rebelein *et al*., 2018). A partial least square discriminant analysis (PLS-DA) was used for separation of groups. Important metabolites were ranked based on Variable Importance in Projection (VIP) score of the PLS-DA. Significantly different metabolites were identified using the Significance Analysis of Microarray (SAM) approach with a Delta of 0.5 within MetaboAnalyst. Metabolites that showed a particular pattern were identified using PatternHunter analysis within MetaboAnalyst (Pavlidis & Noble, 2001).

Unless otherwise indicated, data are shown as means ± standard errors of means (SEM). Differences were considered significant if probability of Type I error was <0.05.

## RESULTS

### Effects of warming and OA on the SMR

Warming significantly elevated SMR (P < 0.001) of *M.edulis* (Fig. 1A, Table 2). After four weeks acclimation, SMR was ~ 2-3 times higher in mussels maintained at 15 °C compared to 10 °C. This effect was observed under normocapnia (P = 0.002) and hypercapnia (P = 0.009). OA did not significantly affect SMR in mussels (P = 0.070), although a trend of elevated SMR was observed in OA-exposed mussels.

**Figure 1:**
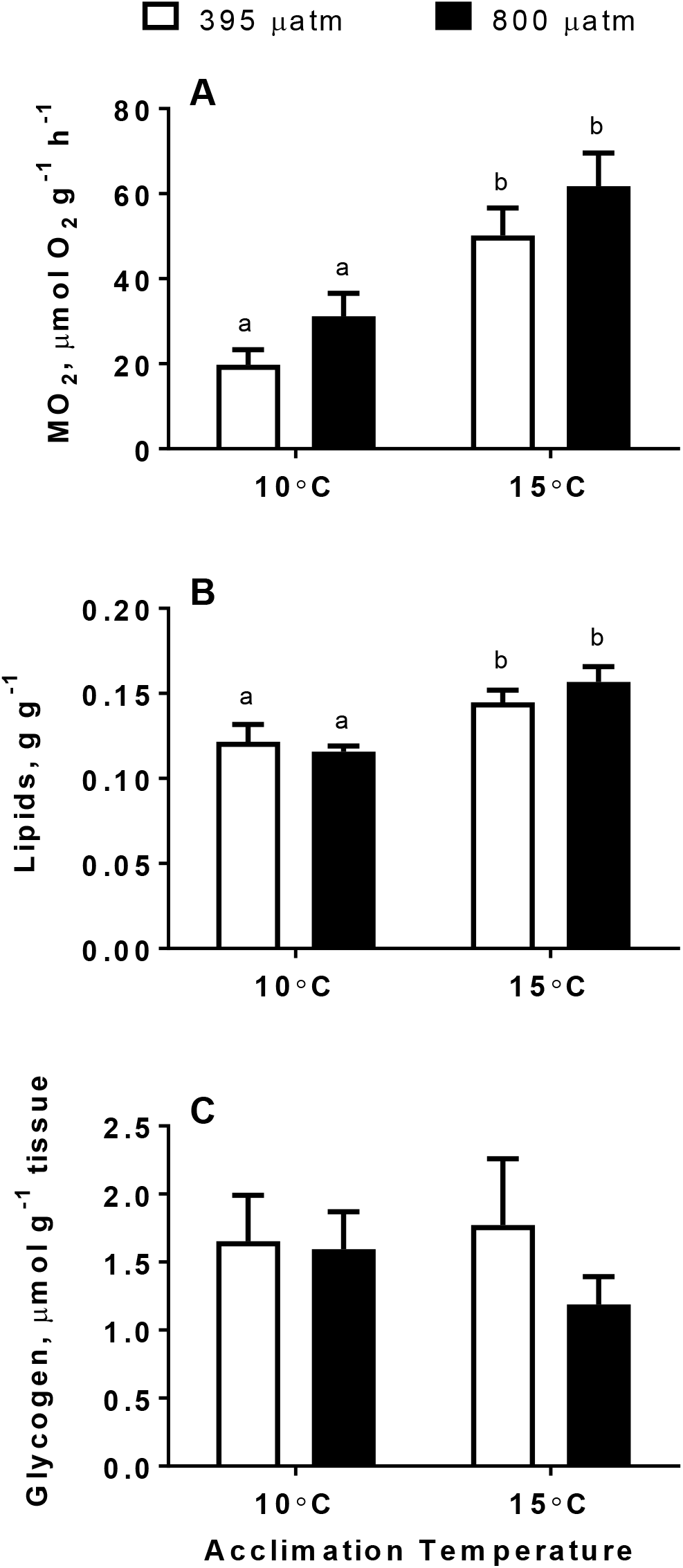
Oxygen consumption rates and tissue energy reserves of M. edulis exposed to different temperatures and P_CO2_. A — oxygen consumption rates (M O2), B — total lipids in hepatopancreas and C — glycogen in adductor muscle. X-axis —temperature. Within each graph, different letters indicate means are significantly different between temperatures within the same P_CO2_ group (P<0.05). Asterisks (*) indicate values that are significantly different between normocapnia and hypercapnia within the same temperature (P<0.05). Vertical bars represent SEM. For M O_2_, N = 8–10. For lipids and glycogen, N = 6–10.

**Table 2:** ANOVA results of the effects of exposure temperature, P_CO2_ and their interaction on energy related indices and enzyme activities in *M.edulis*. F-values are given with degrees of freedom for the factor and error in the subscript. Significant values (P<0.05) are highlighted in bold. AEC, Adenylate energy change.

| Parameter | Temperature | P<sub>CO2</sub> | Temperature X P<sub>CO2</sub> |
| --- | --- | --- | --- |
| Standard Metabolic Rate (SMR) | F<sub>1, 28</sub>=24.97, P< 0.0001 | F<sub>1,28</sub>=3.55, P=0.07 | F<sub>1,28</sub>=0.00, P=0.98 |
| Total Lipids | F<sub>1,20</sub>=16.63, P=0.0006 | F<sub>1,20</sub>=0.16, P=0.69 | F<sub>1,20</sub>=1.30, P=0.26 |
| Glycogen | F<sub>1,35</sub>=0.17, P=0.68 | F<sub>1,35</sub>=0.85, P=0.36 | F<sub>1,35</sub>=0.56, P=0.45 |
| ATP | F<sub>1,20</sub>=0.61, P=0.44 | F<sub>1,20</sub>=3.16, P=0.09 | F<sub>1,20</sub>=0.03, P=0.86 |
| ADP | F<sub>1,20</sub>=3.14, P=0.09 | F<sub>1,20</sub>=0.70, P=0.41 | F<sub>1,20</sub>=0.38, P=0.54 |
| AMP | F<sub>1,20</sub>=1.08, P=0.31 | F<sub>1,20</sub>=0.14, P=0.71 | F<sub>1,20</sub>=0.95, P=0.34 |
| AEC | F<sub>1,20</sub>=2.39, P=0.13 | F<sub>1,20</sub>=0.97, P=0.33 | F<sub>1,20</sub>=0.16, P=0.69 |
| Carbonic Anhydrase | F<sub>1,20</sub>=6.70, P=0.01 | F<sub>1,20</sub>=1.17, P=0.29 | F<sub>1,20</sub>=0.04, P=0.83 |
| Calcium ATPase | F<sub>1,19</sub>=9.86, P=0.005 | F<sub>1,19</sub>=0.01, P=0.90 | F<sub>1,20</sub>=0.18, P=0.67 |
| Proton ATPase | F<sub>1,7</sub>=0.03, P=0.86 | F<sub>1,17</sub>=1.13, P=0.30 | F<sub>1,17</sub>=3.25, P=0.08 |

### Tissue energy status under warming and OA

Warming significantly increased the total lipid content in HP under normocapnia (P = 0.05) and hypercapnia (P = 0.001) (Fig. 1B, Table 2). OA did not significantly change the lipid content in HP, regardless of the temperature (P = 0.608 and 0.288 at 10 and 15 °C, respectively).

No significant changes were observed for glycogen (Fig. 1C) and adenylates (Fig. 2) content under warming, OA, or OA combined with warming (OWA) in the muscle of *M.edulis* (Table 2).

**Figure 2:**
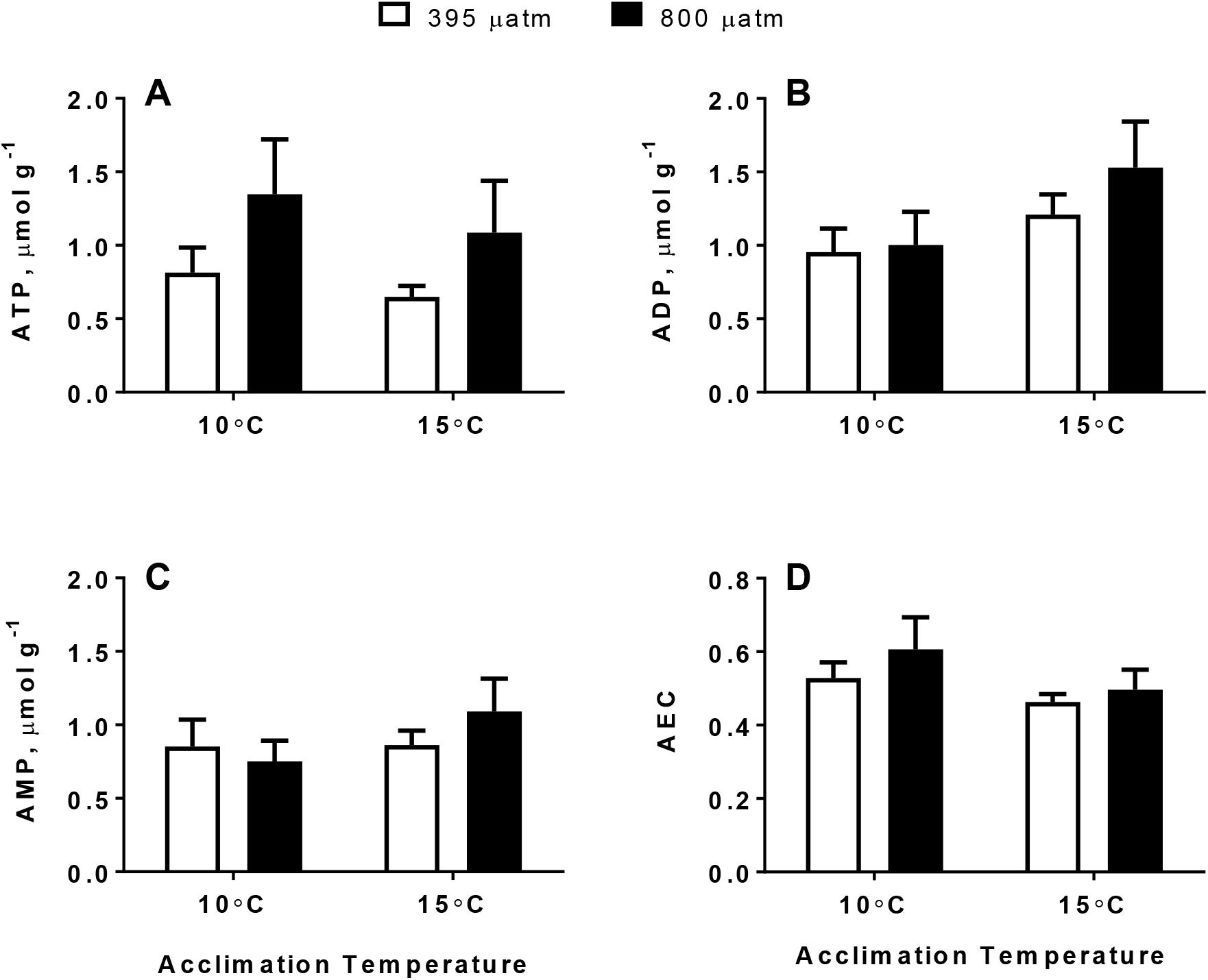
Tissue concentrations of adenylates of *M. edulis* exposed to different temperatures and P_CO2_. A – ATP, B – ADP, C – AMP and D – adenylate energy charge (AEC). If the columns have no letters, the respective means are not significantly different between different P_CO2_ and temperature (P>0.05). Vertical bars represent SEM. N = 6.

### Tissue-specific shifts in metabolite profile under warming and OA

Untargeted NMR-based metabolic profiling in gill and muscle of *M. edulis* revealed minor shifts in metabolite concentrations under warming, OA and OWA. In gills, PLS-DA revealed a significant separation of 10 °C- and 15°C-acclimated groups (Fig. 3A), indicating a temperature-induced change in branchial metabolism. According to their VIP scores, DMA and glucose were the main metabolites describing these differences (Fig. 3B). The levels of DMA decreased and glucose levels increased under warming, regardless of P_CO2_ (Fig. 4A). These two metabolites were also identified as significantly different between the temperature groups by SAM analysis (Fig. 4B). Unlike gills, metabolite profiles of adductor muscle were not affected by temperature or P_CO2_ (see PLS-DA in Supplementary Fig. 3).

**Figure 3:**
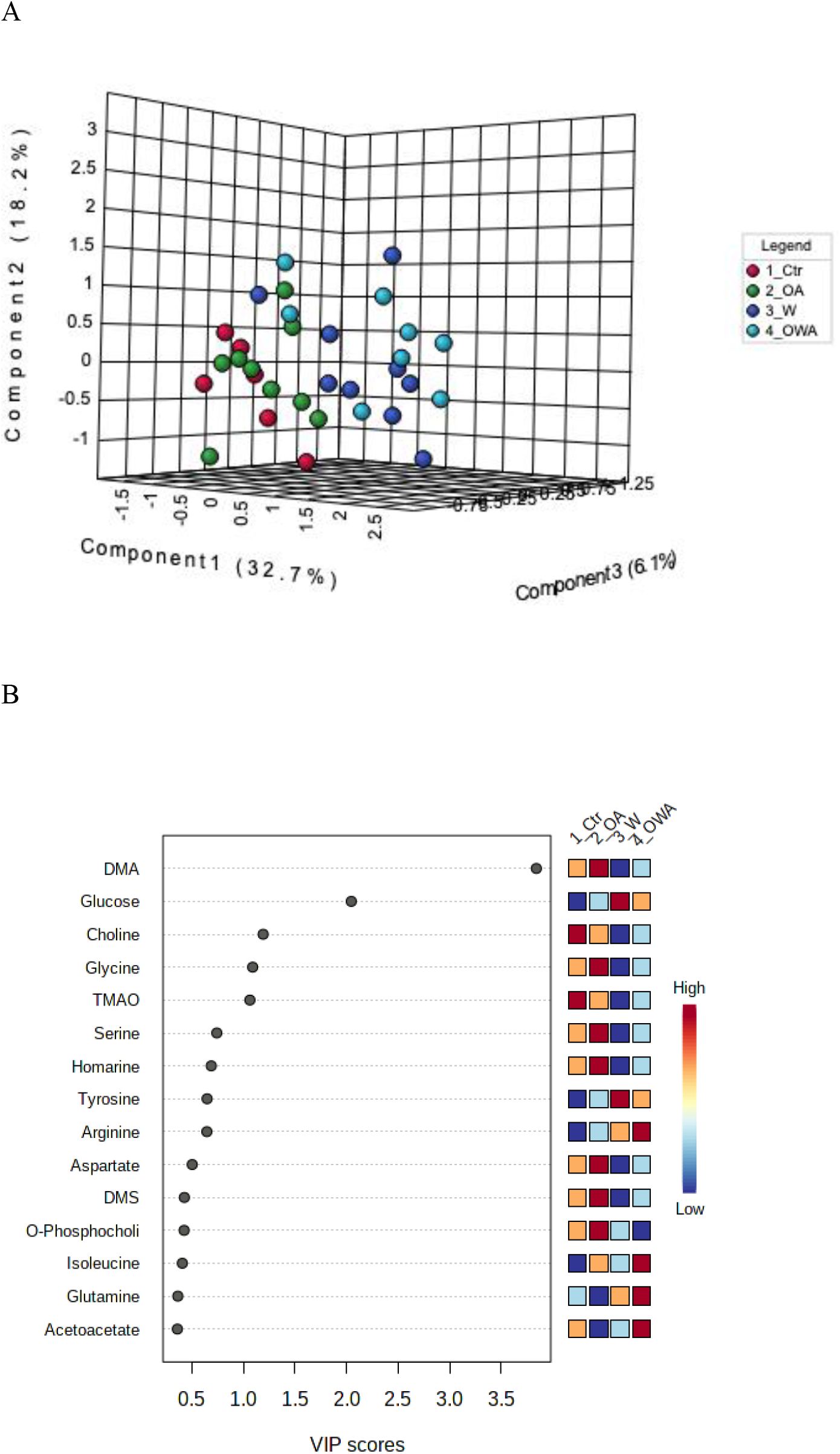
PLS-DA analysis of metabolite profiles in the gill tissues of *M. edulis* exposed to different temperatures and P_CO2_. A – 3D-loading plot of first three components separating the metabolic profiles from Warming and OWA groups (blueish dots) from control (red dots). The OA group (green dots) did not separated from control. B –Important metabolites identified by PLS-DA. The colored squares on the right show group specific relative changes in metabolite concentration. Groups: Ctr – control (acclimated at 10 °C and normocapnia), OA – ocean acidification (acclimated at 10 °C and hypercapnia), W – warming (acclimated at 15°C and normocapnia), OWA – ocean warming and acidification (acclimated at 15 °C and hypercapnia)

**Figure 4A:**
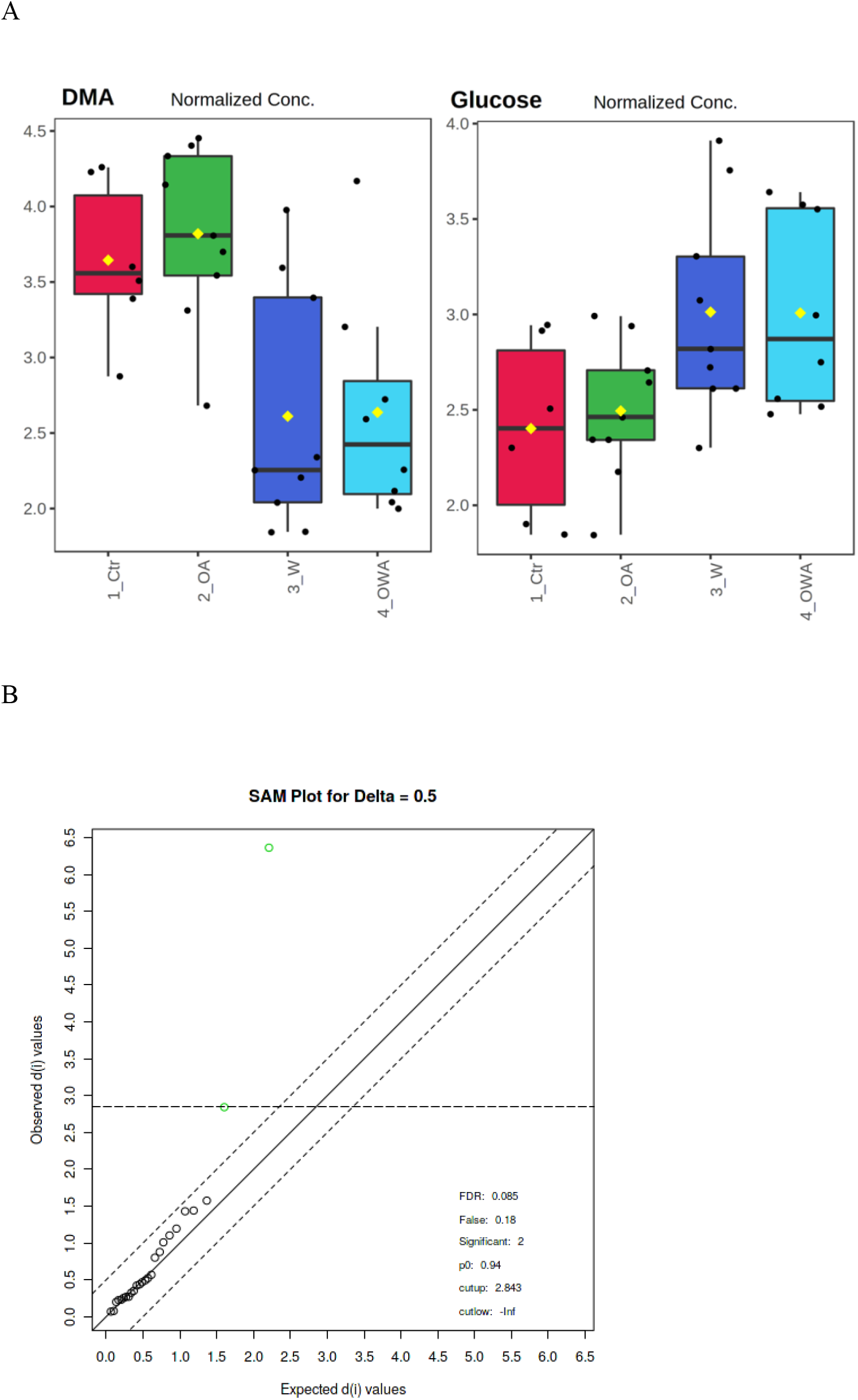
Tissue levels (given in normalized concentrations) of DMA and glucose in gill tissues of *M. edulis* exposed to different temperatures and P_CO2_. Groups: Ctr – control (acclimated at 10 °C and normocapnia), OA – ocean acidification (acclimated at 10 °C and hypercapnia), W – warming (acclimated at 15 °C and normocapnia), OWA – ocean warming and acidification (acclimated at 15 °C and hypercapnia). Figure 4B: Significance Analysis of Microarray (SAM) plot of gill tissue. Scatter plot showing observed relative differences on the axis of ordinates against the expected relative differences estimated by data permutation on the abscise using a delta of 0.5 (dotted lines). The green dots are highlighting significant differences and correspond to DMA and glucose.

### Effects of warming and OA on enzyme activity

CA activity in mantle of *M.edulis* showed an elevated trend under warming (normocapnia, P = 0.093; hypercapnia, P = 0.072) but did not under OA (P = 0.293) (Fig. 5A, Table 2). When data for normocapnia and hypercapnia were pooled, a significant increase in CA activity at 15 °C compared to 10 °C (P = 0.018) was detected (Supplemental Fig. 1). Tissue-specific CA activity over a broad temperature range (5-35 °C) showed a significant effect of two-factor interactions (temperature x tissue, P <0.0001). CA activity was significantly higher in HP compared to other tissues (Fig. 5B, Table 3). Irrespective of the tissue, CA activity monotonously increased with increasing temperatures with similar E_a_, and no ABT (Table 3).

**Figure 5:**
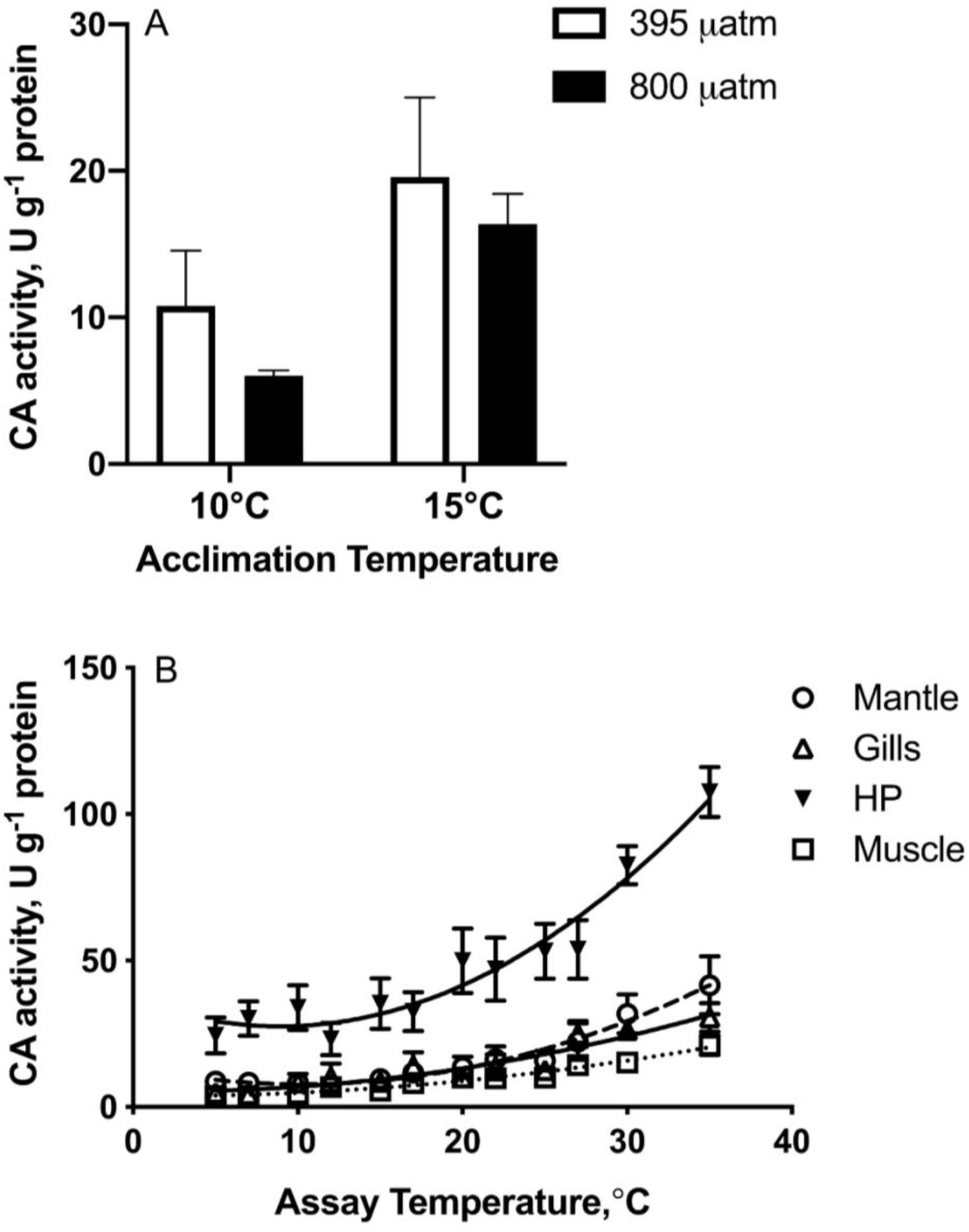
Carbonic anhydrase (CA) activity in tissues of *M.edulis*. A— CA activity in mantle edge exposed to different temperature and P_CO2_. B– tissue-specific variation in specific activities of carbonic anhydrase with temperature. Vertical bars represent standard errors of means. HP-hepatopancreas. If the columns have no letters, the respective means are not significantly different (P>0.05). Vertical bars represent SEM. N= 5–7.

**Table 3:** Activation energies (E_a_) and Arrhenius breakpoint temperature (ABT) for specific activities of carbonic anhydrase in different tissues of *M. edulis*. E_a_ values highlighted in bold and marked with an asterisk are significant after the sequential Bonferroni corrections (P<0.05). Q10 temperature coefficients were calculated for the complete range of the studied temperatures (5-35 °C) and given in the last column.

|  | Carbonic anhydrase |  |  |  |
| --- | --- | --- | --- | --- |
| Tissue | ABT, °C | E<sub>a</sub>, kJ mol<sup>-1</sup> K<sup>-1</sup> |  | Q<sub>10</sub> |
|  |  | Below<br>ABT | Above<br>ABT |  |
| *M. edulis* |  |  |  |  |
| Mantle | N/a | 39.82* | - | 1.68 |
| Gills | N/a | 43.81* | - | 1.84 |
| Hepatopancreas | N/a | 32.50* | - | 1.64 |
| Muscle | N/a | 40.98* | - | 1.77 |

Ca^2+^-ATPase activity from mantle edge of *M.edulis* was significantly affected by warming (P = 0.005) but not OA (P = 0.905) (Fig. 6A, Table 2). Warming led to a significant decrease in Ca^2+^-ATPase activity under hypercapnia (P = 0.023) but not under normocapnia (P = 0.060).

**Figure 6:**
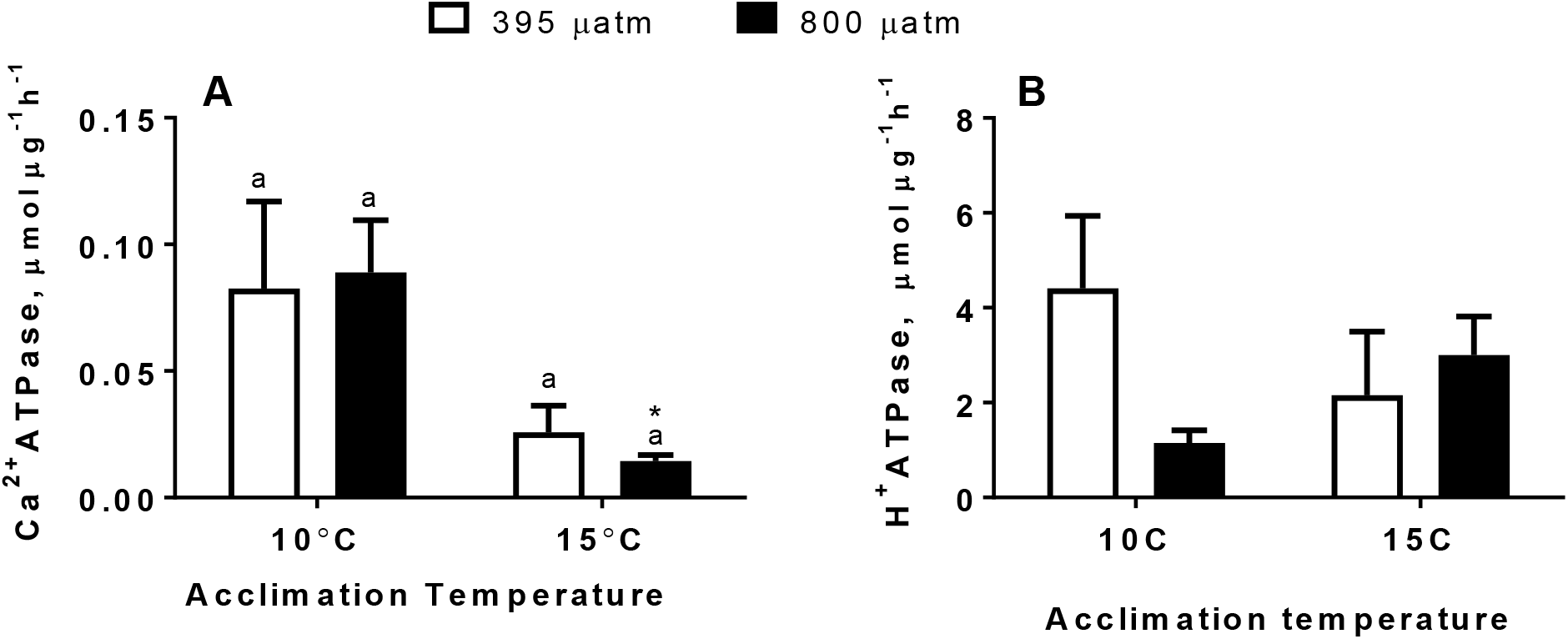
ATPases activity in mantle edge of *M.edulis* exposed to different temperatures and P_CO2_ levels. A – calcium (Ca^2+^) ATPase and B – proton (H^+^) ATPase. If the columns share letters or have no letters, the respective means are not significantly different (P>0.05). Asterisks (*) indicate values that are significantly different between normocapnia and hypercapnia within the same temperature (P<0.05). Vertical bars represent SEM. N= 5-6.

H^+^-ATPase activity from mantle edge of *M.edulis* was not significantly affected by warming (P = 0.862) or OA (P = 0.303) (Fig. 6B, Table 2). At 10 °C, hypercapnia led to a decrease in the H^+^-ATPase activity, but this decrease was non-significant (P = 0.06).

## DISCUSSION

Our study demonstrates that adult *M.edulis* from GOM are metabolically resilient to moderate OA (P_CO2_ ~800 μatm) but responsive to warming as seen in changes in whole-body metabolic rate, energy reserves, metabolite profiles and enzyme activities. The combination of ocean warming and acidification (OWA) did not elicit detrimental metabolic changes in mussels beyond the effect of warming. This indicates that under these conditions temperature is the dominant factor in determining species’ physiology. The physiological responses of mussels in our study might be related to their ecology in GOM. There is a high temporal variability in temperature (annual SST range 15.5 °C) in temperate rocky intertidal pools of GOM throughout the year (Salisbury & Jönsson, 2018). Modelling data indicate that physical processes (e.g., strong tides, wind-driven mixing) in GOM alter ocean carbonate parameters but also mitigate the decrease, or even raise pH (Salisbury and Jönsson 2018). Thus, *M. edulis* in this region are predominantly exposed to large variation in temperature which might explain their plasticity to thermal stress. Earlier studies in *M.edulis* populations from thermal clines of GOM have shown high phenotypic plasticity in physiology despite lack of population genetic structure and local adaptation (Lesser *et al*., 2010; Lesser, 2016) consistent with the findings of metabolic plasticity to warming found in our study.

### Effect of Warming and OA on Bioenergetics

We observed a ~2-3-fold increase in SMR of mussels exposed to 5 °C warming under normocapnia or hypercapnia, indicating a strong temperature effect on metabolism with Q10 ~4-6. A 5 °C increase is well within the range of temperature fluctuations experienced by mussels in GOM (Salisbury & Jönsson, 2018). Blue mussels are eurythermal and well adapted for 5-20 °C range, with an upper thermal tolerance limit of ~29 °C for adults (Gosling, 1992). Therefore, temperatures in our study are within the ecological relevant and even optimal range for this species. Recent meta-analysis also indicates that temperature threshold for long-term survival of *M. edulis* is ~20 °C (Lupo *et al*., 2020). Therefore, 20 °C appears to be close to the metabolic optimum of *M. edulis*, so rate-enhancing effects of temperature dominate over the potentially negative impacts on metabolism as long as warming occurs below the 20 °C threshold, as in our study.

Unlike temperature effect, modest OA had no effect on SMR regardless of the temperature. Metabolic response to OA in *M. edulis* can vary and depend on magnitude of P_CO2_, food availability, and population (Fitzer *et al*., 2014; Hüning *et al*., 2013; Lesser, 2016; Thomsen *et al*., 2013; Thomsen & Melzner, 2010; Zittier *et al*., 2015). Fitzer *et al*. (2014) have shown that for *M.edulis* at 1000 μatm and beyond, biomineralization continued but with compensated protein metabolism and shell growth indicating that ~1000 μatm could be an OA metabolic tipping point for *M.edulis*. In our study, P_CO2_ levels were below this threshold and could explain the physiological tolerance of mussels seen here. Our hypothesis that P_CO2_ could exacerbate the increase of SMR caused by elevated temperature was not supported in this study. The outcome of OWA on metabolic rate in bivalves is commonly additive; albeit in other cases the effects of temperature or P_CO2_ dominate (see Lefevre 2016 and references therein). Furthermore, metabolic responses of bivalves to OWA are dependent on the degree of temperature or P_CO2_ stress. For example, Lesser *et al*., 2016 have shown that mussels from GOM showed metabolic depression as a protective response when exposed to combined stress of higher warming (22 °C) and modestly elevated P_CO2_ (560 μatm).

Temperature (but not OA) had a marked effect on energy reserves in *M. edulis*. We observed an increase in lipids under warming in HP, under both normocapnia and hypercapnia. In bivalves, lipids are primarily stored in HP (Giese, 1966) and synthesis, storage and use of lipids show pronounced seasonal cycles: lipids are accumulated during summer (at high temperature and food availability) and used for metabolism and initiation of gametogenesis during winter (at low temperature and food availability) (Fokina *et al*., 2015). Laboratory studies indicate that lipid accumulation is a direct response to temperature in mussels (Fokina *et al*., 2015; Wu *et al*., 2021) and might reflect a metabolic adjustment for anticipated reproduction (which requires high energy investment as well as lipid deposition into developing gametes) in mussels.

Unlike lipids, the glycogen store was unresponsive to warming in mussels from GOM. Baltic Sea mussels also showed no change in glycogen content during warming from 10 to 15 °C and from 15 to 20 °C (Wu *et al*., 2021). Modest hypercapnia likewise had no effect on the glycogen content of adductor muscle of mussels in our present study. Earlier studies show that impacts of temperature and OA on glycogen reserves of mussels are threshold dependent. Thus, mussels from GOM showed a marked decrease in the glycogen content of adductor muscle when exposed to higher temperature (22 °C) alone and combined with modest acidification (560 μatm P_CO2_, pH 7.9) (Lesser *et al*., 2016). The concentrations of adenylates and AEC in adductor muscle remained at steady-state levels in all exposures, indicating that cellular energy balance in mussels was maintained under all temperatures and OA scenarios. Overall, our findings indicate that temperature and OA used in our present study are not energetically stressful to GOM mussels (at least under ad libitum feeding conditions).

Warming alone or OWA altered the metabolite profile of *M. edulis* in a tissue-dependent manner. Warming from 10 to 15 °C increased glucose and decreased DMA in the gills, whereas adductor muscle metabolites were not affected. Increased glucose could reflect mobilization of energy reserves to meet increased tissue energy demand. Furthermore, warming could increase glucose levels via gluconeogenesis by channeling of amino acids like serine, alanine and glycine into pyruvate and increased activity of the enzyme phosphoenolpyruvate carboxykinase (PEPCK) (Ellis *et al*., 2014; Le Moullac *et al*., 2007). In our study, we saw a trend for decreased glycine and serine in gills under warming and OWA, suggesting a potential for increased flux through gluconeogenesis that warrants further investigation. The decrease in DMA, a common organic osmolyte found in gills of bivalves (Zhang *et al*., 2011), might osmotically compensate for elevated glucose in *M. edulis* gills during warming.

In *M. edulis*, OA had no effect on the metabolite profile in gills or muscle tissues. Similarly, metabolite profiling studies with P_CO2_ ≤ ~1,000 μatm (pH ≥ ~7.8) reported no OA-induced alteration in metabolite levels of bivalves (Dickinson *et al*., 2012; Ellis *et al*., 2014; Wei *et al*., 2015) whereas higher P_CO2_ (≥1,500 μatm, pH ≤ ~7.7) led to a shift in metabolite profiles with alterations in energy metabolism (Ellis *et al*., 2014; Lannig *et al*., 2010; Wei *et al*., 2015). These findings are consistent with the notion that modest OA is not a metabolic stressor for *M. edulis*, and the studied GOM population follows this general pattern.

### Effect of Warming and OA on Enzyme Activity

In mollusks, CA plays a key role in the maintenance of acid-base homeostasis of all tissues as well as biomineralization in the mantle (Li *et al*., 2016; Wang *et al*., 2017; Ramesh et al., 2020). In *M. edulis*, CA activity increased with acute warming (5-35 °C) in all tissues with similar E_a_ values indicating a concerted whole-body response of this enzyme to warming. No ABT was found for CA activity from 5 to 35 °C indicating high thermal tolerance of this enzyme. We found highest CA levels in HP as compared to other tissues (gills, muscle and mantle) in *M. edulis*. This might reflect differences in overall metabolic activity (and therefore, different metabolic CO_2_ and proton loads) among the tissues that require different levels of CA to maintain acid-base balance. Acclimation to 15 °C upregulated CA activity in a key biomineralizing tissue (the mantle edge) of *M. edulis*. Similarly, long term (15 weeks) exposure to elevated temperature (27 °C) led to a notable increase in CA activity in bivalves *Crassostrea virginica* and *Mercenaria mercenaria* (Ivanina *et al*., 2013). This increase is likely linked with the overall increase in metabolic rates at elevated temperatures and can assist with shell deposition.

CA activity remained unchanged in response to OA in mantle edge of *M. edulis*. Previous studies have shown variable CA responses to OA in bivalves including mussels (Ivanina *et al*., 2020). Similar to our findings, CA activity in the mantle of *C.virginica* and *M. mercenaria* remained unchanged after exposure to elevated P_CO2_ (800 μatm) for 2-15 weeks (Ivanina *et al*., 2013). In contrast, CA activity in mantle of *M.edulis* decreased after prolonged (6-months) exposure to elevated P_CO2_ (750 μatm) exposure (Fitzer *et al*., 2014). In oysters, CA accumulated along mantle edge in response to P_CO2_ exposure (2622 μatm, pH 7.50), suggesting an active role of CA in ion-regulation and acid-base balance (Wang *et al*., 2017). However, the effect of OA on CA activity is threshold dependent; at P_CO2_ <1000 μatm mussels try to compensate for intracellular acid loads instead of decreasing their metabolism (Fitzer *et al*., 2014; Hüning et al., 2013; Thomsen & Melzner, 2010). Taken together, these data suggest that *M. edulis* in GOM can upregulate acid-base capacity contributing to their metabolic plasticity towards warming, but moderate OA has no effect on this trait.

As a consequence of maintaining acid-base balance, OA may change concentrations of H^+^, HCO_3_^-^ Ca^2+^, Mg^2+^, and Cl^-^ in calcifiers (Fitzer *et al*., 2014; Ramesh *et al*., 2017). Ion transport is an important contributor to energy budget of biomineralization because Ca^2+^ transport and removal of excess protons from the site of biomineralization are ATP-dependent (Ivanina *et al*., 2020). In GOM mussels, activity of H^+^-ATPase and Ca^2+^-ATPase-in the mantle remained stable under moderate warming and OA (except for a modest but significant decline in Ca^2+^-ATPase under OWA). This indicates that the mussels can maintain ion regulatory fluxes at least in the mantle edge despite variations in temperature and P_CO2_ relevant to near-future climate change. Mussels from other acidified environments like Kiel Fjord also build and maintain their shells despite fluctuating P_CO2_ and pH, partially owing to enhanced ion transport (Thomsen *et al*., 2013, 2017). Furthermore, when mussel larvae were raised under OA between 500-1500 μatm P_CO2_, ΔH^+^ between calcification site and sea water remained constant, irrespective of P_CO2_ (Thomsen *et al*., 2013). In *C. virginica* and*M. mercenaria* activities of Ca^2+^ ATPase and H^+^ ATPase, as well as the cellular energy costs of Ca^2+^ and H^+^ transport in the biomineralizing cells (mantle and hemocytes) were insensitive to ocean acidification (pH 7.8) (Ivanina *et al*., 2020). This indicates that intertidal species (such as mussels, oysters and clams) that are adapted to variable temperature and pH in their habitat are generally tolerant against moderate warming and OA, predicted by the climate change models.

## CONCLUSIONS

In this study, we report that adult *M. edulis* from GOM are sensitive to warming but tolerant to moderate acidification scenario predicted by IPCC for the year 2100. This result also provides an insight in the natural history of GOM mussels given that in the last decade (2005-2014) GOM was characterized by an extreme warming trend (Salisbury & Jönsson, 2018). Although this study is limited to adults and does not consider larval stage sensitivity of *M. edulis*, our results support earlier reports that acidification scenarios for the next 100-300 years do not affect this species (Telesca *et al*., 2019). Taken together, our study provides important data about extant levels of plasticity in physiology of mussels as well as insights into potential sensitivity of mussels to future global change.

## Acknowledgements

The authors thanks Dr. Markus Frederich (University of New England) for assistance with mussel collection in Gulf of Maine. The authors declare that there is no conflict of interest. This work was supported by the US National Science Foundation (NSF) award IOS-1557870 to I.M.S.

## Author contributions

Omera B. Matoo and Inna M. Sokolova conceived the ideas and designed methodology; Omera B. Matoo, Gisela Lannig and Christian Bock collected the data; Omera B. Matoo, Gisela Lannig and Christian Bock analyzed the data; Omera B. Matoo, Gisela Lannig, Christian Bock and Inna M. Sokolova led the writing of the manuscript. All authors contributed critically to the drafts and gave final approval for publication.

## Data availability statement

Supplemental files are available at FigShare. Phenotypic data will be deposited in the Dryad Digital Repository upon publication.

