## Supplemental Figures for "Temperature but not ocean acidification affects energy metabolism and enzyme activities in the blue mussel, *Mytilus edulis*"

**Supplemental Figure 1: Carbonic anhydrase activity in mantle edge tissue of blue mussel (*M.edulis*) at exposure temperatures**. Samples from normocapnia (395 µatm) and hypercapnia (800 µatm) were pooled to determine the effect of temperature. Letters indicate respective group means are significantly different from each other (P<0.05). vertical bars represent standard error of means. N= 10-14.

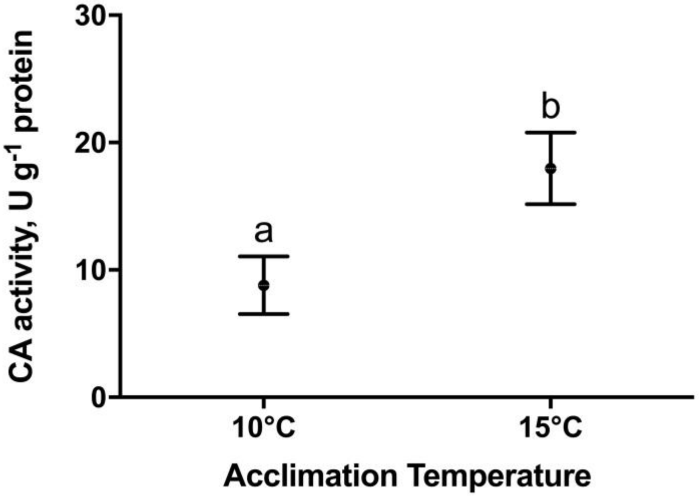

**Supplemental Figure 2: Correlations between CA and total esterase activity levels measured in different tissues of blue mussel (*M.edulis*).** Data points on the regression lines correspond to CA and total esterase activities measured at different temperatures. The slopes are significant (P<0.05) in all tissues.

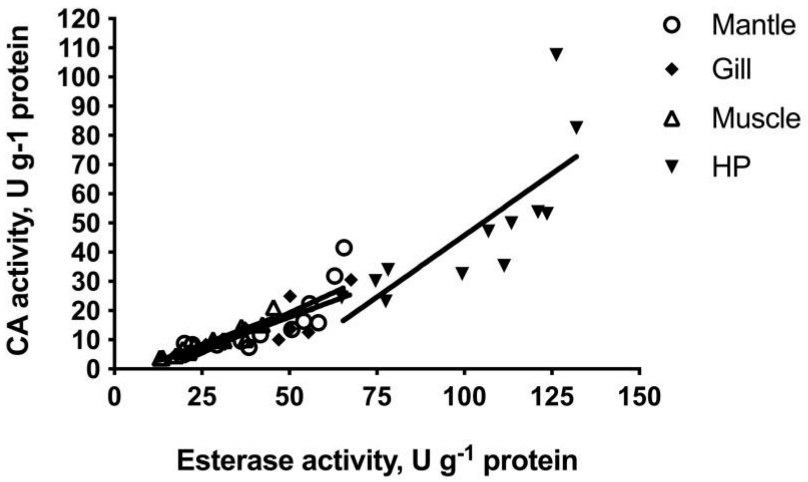

**Supplemental Figure 3: PLS-DA analysis of metabolite profiles in the adductor tissue of M. edulis exposed to different temperatures and $P_{CO2}$.** 3D-loading plot of first three components of the metabolic profiles in Warming and OWA groups (blueish dots), OA group (green dots) from control (red dots). There is no separation in the metabolic profiles under any treatment.

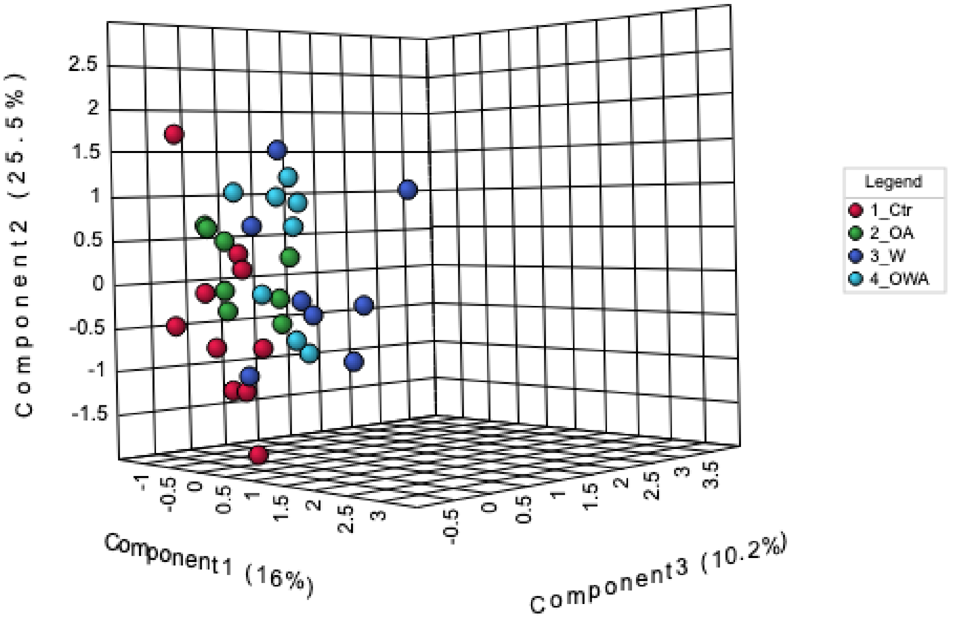
